# Eukaryotic chemotaxis under periodic stimulation shows temporal gradient dependence

**DOI:** 10.1101/2023.10.17.562804

**Authors:** Richa Karmakar, Aravind Karanam, Man-Ho Tang, Wouter-Jan Rappel

## Abstract

When cells of the social amoeba *Dictyostelium discoideum* are starved of nutrients they start to synthesize and secrete the chemical messenger and chemoattractant cyclic Adenosine Mono Phosphate (cAMP). This signal is relayed by other cells, resulting in the establishment of periodic waves. The cells aggregate through chemotaxis towards the center of these waves. We investigated the chemotactic response of individual cells to repeated exposure to waves of cAMP generated by a microfluidic device. When the period of the waves is short, the chemotactic ability of the cells was found to increase upon exposure to more waves, suggesting the development of a longer-term memory. This effect was not significant for longer wave periods. We show that the experimental results are consistent with a model that includes a slowly rising and decaying component that is activated by the temporal gradient of cAMP concentration. The observed enhancement in chemotaxis is relevant to populations in the wild: once sustained, periodic waves of the chemoattractant are established, it is beneficial to cells to improve their chemotactic ability in order to reach the aggregation center sooner.

---

Chemotaxis, the chemically guided motion of cells, is critical to several biological processes such as foraging, wound healing, embryonic developement, and cancer metastasis [1–5]. The social amoeba *Dictyostelium discoideum* is a well-characterized model organism to study chemotaxis. It displays a unicellular to multicellular transition in its life cycle when starved of nutrients, by aggregating through chemotaxis [6, 7]. Chemotaxis of *Dictyostelium* cells is driven by the chemoattractant cyclic Adenosine Mono Phosphate (cAMP), a small molecule that is internally synthesized and secreted by the cells [8]. The multicellular aggregate then forms a stalk and a fruiting body, which contains spores that can later be dispersed [9, 10].

During the aggregation process, *Dictyostelium* cells not only produce the chemoattractant but also relay the signal, resulting in cAMP waves that sweep through the population [11]. This signal-relay process is also present in other chemotactic systems [12, 13] and ensures the recruitment of cells over large distances. Initially, the cAMP waves arise spontaneously from many locations in the population. As development continues, sources with high frequencies dominate and become stable centers to which cells aggregate [14]. A large number of models of the aggregation process have been proposed, ranging from qualitative excitable system models to more biochemically oriented ones [15–19]. In addition, a significant number of modeling studies have been published which attempt to describe the chemotactic response of a single cell to chemoattractant gradients [20–22].

Traditionally, most experimental and modeling chemotaxis studies have focused on the response of cells to static gradients [23, 24]. More recently, using microfluidic devices, it has become possible to expose cells to carefully controlled complex and time-varying gradients [25–30]. For example, experiments that generated chemoattractant waves of controlled speed have shown that cells can be more sensitive to the positive gradient in the incoming half of a wave and less sensitive to the negative gradient in the back half of the wave [31, 32]. The extent of this bias of cell motion has been shown to depend on the period of the chemoattractant waves [31, 32] as well as the background concentration of the chemoattractant [30]. These studies also support the involvement of a Local Excitation Global Inhibition (LEGI) module in chemotaxis [31, 32]. In this module, the stimulus produces both a membrane bound localized activator and a globally diffusible inhibitor, with fast and slow kinetics, respectively [20, 33].

These previous studies did not determine whether cell motion depends on temporal sensing. Furthermore, it did not address the question of the exposure to multiple waves, a question relevant given the aggregation process of *Dictyostelium*. In this study, we investigate the effect of exposure to multiple waves of identical amplitude and frequency on the chemotactic ability of *Dictyostelium* cells. Our results show that this ability is enhanced upon exposure to multiple waves of a short period (6 min) and remains low and unchanged when the period is long (15 and 20 min). Furthermore, using modeling, we show that our result supports a mechanism for temporal gradient sensing and rectification, working in parallel with a mechanism for spatial gradient sensing in the form of a LEGI model, and explains the observed trend in the chemotactic ability of the cells.

We exposed cells of the axenic strain AX4 [34] that are developed for 5 hours to multiple identical cycles of cAMP waves with a uniform speed. These traveling waves were generated using a microfluidic device, detailed in an earlier study [30]. In short, a stream of cAMP is swept across an observation channel, resulting in a bell-shaped wave profile that is similar to the one measured for natural waves of cAMP [15, 35]. The speed of the wave can be controlled such that each wave sweeps through the substrate in a fixed time period *T*. The period was set to either 6 minutes (hereafter referred to as short period), corresponding to a physiologically relevant value [36], or to higher values, namely 15 and 20 minutes (hereafter referred to as long periods). Cells were plated on a micropatterned substrate consisting of one dimensional patterns, which was also used in an earlier work [37]. As a result of this patterning, cells were confined to move along only one dimension, either along or opposite the direction of wave propagation, facilitating cell tracking. For each wave period, we carried out experiments on at least three different days.

The motion of the cells was captured by Differential Interference Contrast (DIC) microscopy. The cells in the resulting images were smoothened, segmented through edge detection, and binarized. Then, cell tracks were constructed by identifying the nearest neighbor for each cell in the subsequent frame. Only isolated cells were considered for tracking to preclude the effects of crowding and cell contacts on chemotaxis. Furthermore, we only analyzed cells that were tracked for at least two successive waves. The total number of cells tracked for each cycle and period ranged from 125 to 648. To keep the total time experimentation time equivalent, we exposed cells to 7, 4, and 3 cycles of *T* = 6, 15, and 20 minutes, respectively.

Once the tracks of isolated cells were constructed, we computed a measure of the chemotactic ability of the cells called the Chemotactic Index (CI), defined as the ratio of the velocity of the cell in the direction of the source to the magnitude of its speed, averaged over a moving time-window of a certain length (2*τ*) [31]. It follows, then, that CI takes on values between −1 (motion exactly away from the source) and +1 (motion exactly towards the source). In the current one-dimensional context, where cells move only in the *x*-direction, the CI of a cell at time *t* is the ratio of its displacement to its path length over the duration 2*τ*.

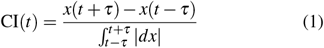

In this study, we have taken the moving window duration 2*τ* to be 2 minutes. After obtaining the time-series, we computed the average chemotactic index, ⟨CI⟩, for each cycle, which allowed us to determine the dependence of ⟨CI⟩ on the number of waves experienced by the cells.

Figure 1-a shows the CI vs time data for all the wave periods and cycles used in the study. For the *T* = 6 minute waves, the CI increases uniformly during the first few cycles, and then saturates. In contrast, the CI for the longer wave periods remains almost unchanged for each cycle. This is also evident in Figure 1-b, which plots the average CI as a function of the wave cycle. From this figure, it is clear that ⟨CI⟩ increases substantially (by about 50%) for *T* = 6 min and it remains roughly the same for the longer wave periods. We have verified that the increase in CI is due to an increase in the directionality of the cell motion: the (undirected) cell speed is roughly the same (∼3 −4 μmmin^−1^) for all cycles and periods (see Supplemental Material Figure 1).

**FIG. 1.**
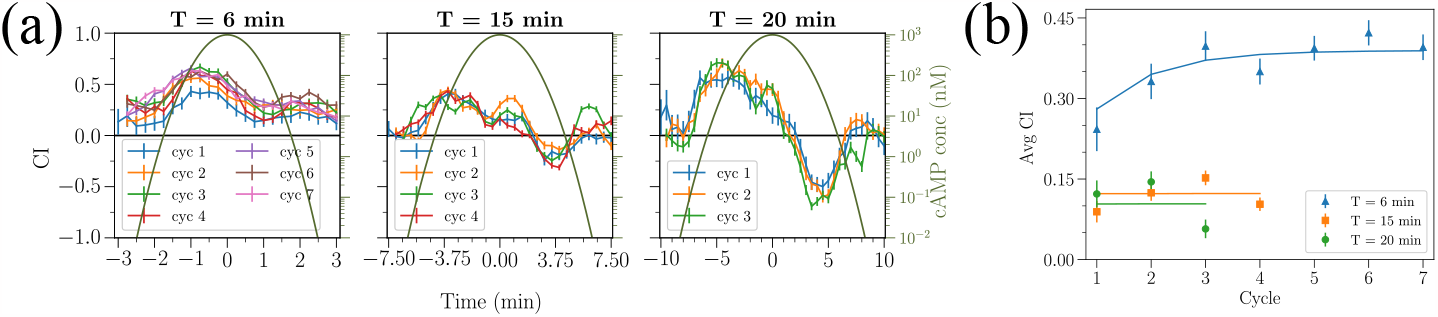
(a) Chemotactic Index (CI) as a function of time for time periods *T* = 6 min, 15 min, and 20 min, each with several cycles. Each data point represents the average CI of all the cells in the time bin-width Δ*t*. Error bars represent the standard error of the mean. Δ*t* is taken to be the same as the frame rate of the image capture, i.e., 15 s for *T* = 6 min and 30 s for *T* = 15 min and 20 min. Also shown is the cAMP profile concentration for each wave period. (b) Average CI plotted against cycle number for all periods. Markers represent an average over all tracked cells and error bars represent the standard error of the mean. Lines correspond to the fits from the model.

To determine whether the increase in the chemotactic ability was due to an increase in development time, we repeated the experiments using cells that were developed for only four hours. These cells also exhibited a significant increase in ⟨CI⟩ for *T* = 6 min waves, similar to 5 h developed cells (see Supplemental Material Fig. 2). We should note that carrying out experiments for cells that were developed for 6 hours or more was challenging since these cells displayed increased adhesion and started to clump into small aggregates. In summary, our experiments show that for short periods, the chemotactic ability of cells markedly improved as the number of waves increased, independent of the development time. In contrast, for longer wave periods, the chemotactic ability was found to be independent of the wave cycle.

**FIG. 2.**
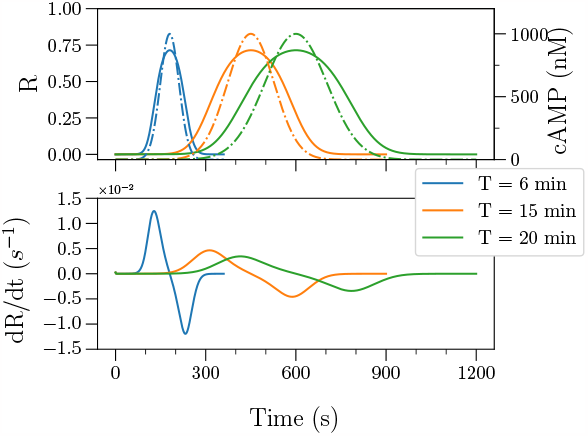
Variation with time of the concentration and temporal gradient of *R* for different periods, for one cycle. cAMP concentration is shown in the top panel with dashed lines. For all wave periods, cAMP and *R* have the same amplitude, but since the wave takes different times to sweep through the substrate, their temporal gradients are different. A wave with a short period has a higher temporal gradient, as shown in the bottom panel. This plot shows the values at the front of the cell; those at the back take on similar values with a small time lag.

We next attempted to address our experimental findings within a modeling framework. The starting point for this is our previous model, which consists of a LEGI module, together with a bistable memory module *M* [31] (see Figure 3-a). In this LEGI+M model, external cAMP binds to the receptor *R*, which then activates both a membrane-bound activator, *E*, and a global inhibitor *I*. The membrane-bound response element *S* of the LEGI module is activated by *E* and inhibited by *I* and feeds into the memory module. The component of this module feeds back to *S* and this feedback depends on *E*. This model was able to explain how cells are able to chemotax toward the wave source, even though the spatial gradient reverses direction in the back of the wave. It was shown that for short period waves the memory at the front, but not the back, is activated, resulting in a continued response in the direction of the original wave [31]. Specifically, the CI was assumed to be determined by a linear combination of the difference in front and back value of *M* and *S*. Importantly, in this study, the parameter values of the LEGI and M modules are identical to the ones used in our previous study and are listed in Supplementary Material Table 1.

**FIG. 3.**
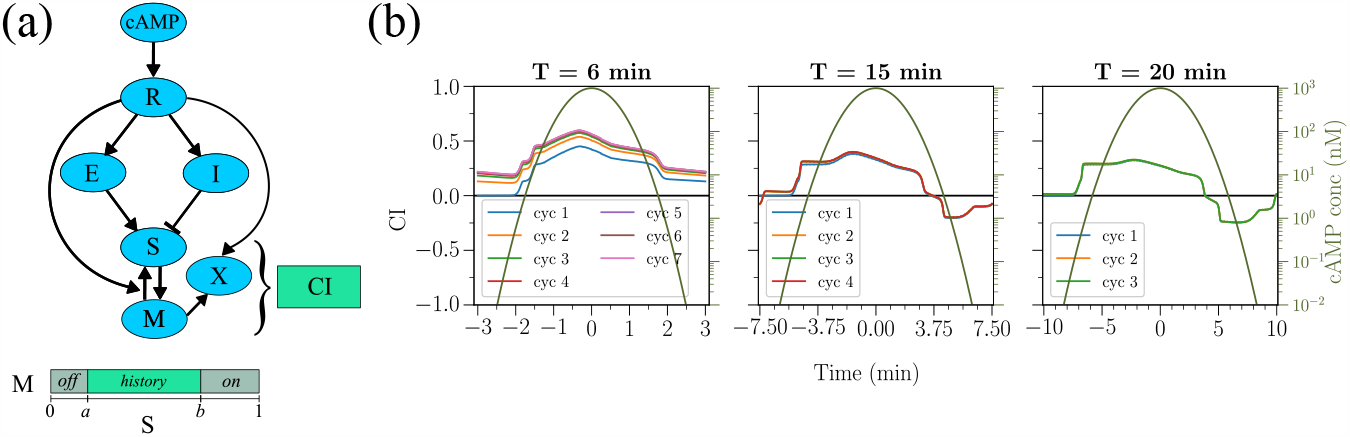
(a) Schematic of the model, by extending the LEGI+M model to include *X* that is activated by the temporal gradient of cAMP with a threshold. See Section 4 in the SI for equations. (b) CI vs time for multiple periods and all cycles as predicted from the model (*α* = *β* = *γ* = 0.15). The plots overlap for *T* = 15 min and 20 min.

**TABLE 1:**
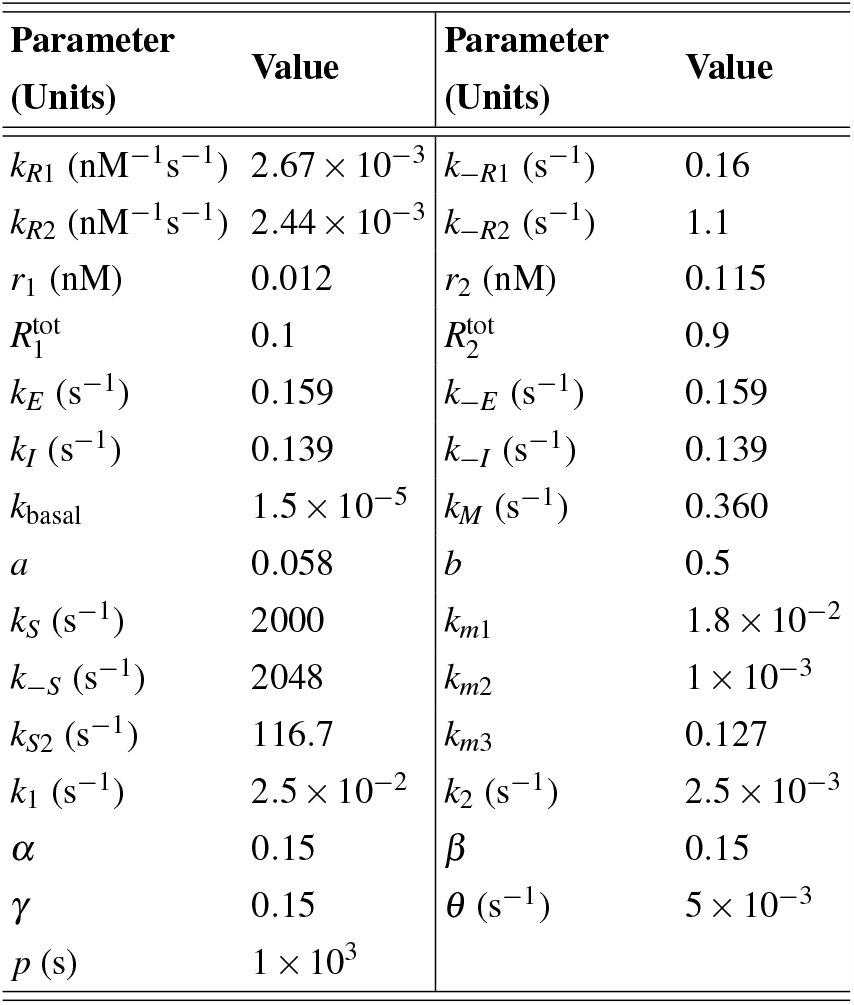
Parameters of the extended LEGI+MX model. The values of *M*^tot^, *S*^tot^ are set to 1 at the front as well as at the back.

Here we extend the model with a new, local signaling component, *X*, whose dynamics is assumed to be a function of the fraction of bound receptors *R* and the memory module *M*. (Figure 3-a, see Supplemental Material Section IV for explicit equations). The model is implemented in a one-dimensional geometry, consisting of a line, representing the interior of the cell, and the two ends corresponding to the back (b) and front (f) of the cell. Thus, all components, including *X*, are solved for both at the front and at the back, except for the global inhibitor *I*. This global component diffuses within the interior of the cell and we will assume that the diffusion coefficient for *I* is large, so that its concentration can be taken as uniform within the cell. Finally, the definition of CI is extended to include a contribution from *X* in addition to *M* and *S*: CI = *α*(*M*_*f*_ − *M*_*b*_) + *β* (*S*_*f*_ − *S*_*b*_) + *γ*(*X*_*f*_ − *X*_*b*_), where *α, β*, and *γ* are parameters that were adjusted to fit the experimental data.

We propose the kinetics of *X* in order to increase the chemotactic response for cells that are exposed to multiple short period waves but not for long period waves. This will be accomplished if *X*_*f*_ − *X*_*b*_ grows as a function of cycle number only for *T* = 6 minute waves. A simple way to arrest the growth of *X* for large wave periods is to introduce a time scale for the decay of *X* that is larger than the short period but smaller than the long period. This would readily work if the activation terms for *X* for both short and long period waves lead to similar increases in the level of *X* after each cycle. A closer look at the wave profile reveals that such an activation cannot depend on the magnitude of the cAMP concentration or any other quantity that directly depends on it, such as the receptor fraction *R*. Figure 2 (top panel) shows the concentration of cAMP (dashed lines) and *R* (bold lines) a cell experiences as a function of time for all three wave periods. Since the difference between the short and long period waves is the speed with which the wave moves over the cells, the profiles of cAMP and *R* have the same amplitude but different widths (in time). Thus, cells exposed to long period waves experience high values of cAMP concentration for a prolonged period of time. Therefore, if the activation of *X* is proportional to the cAMP concentration or depend on a threshold value, than it will reach higher values for long periods than for short periods. Since the chemotactic response is proportional to *X*, this increase will then lead to an elevated CI, in disagreement with our experimental findings. Further details of models that fail to predict the observed trend are described in Section IV of the Supplementary Material.

Cells sense the cAMP concentration through receptor binding, and examining the profiles of *R* experienced by cells reveals that the temporal gradient of *R* is greater for a short wave period than for a long period (see Figure 2, bottom panel). This suggests that the response of cells may involve the temporal gradient of the *R*. Therefore, we propose that *X* is activated only by a temporally increasing cAMP. Specifically, we assume that the dynamics of *X* at the front of the cell can be written as

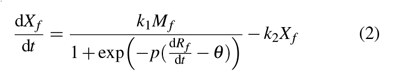

with a similar equation for *X*_*b*_, the component at the back. Here, *M, X*, and *R* are the respective de-dimensionalized concentrations of the respective components in the model. They take values between 0 and 1 at the front as well as at the back. The first term describes the activation of *X* through a sigmoidal function of the temporal concentration 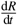 of the activated receptor with a positive threshold *θ*, which acts as a rectifier and places a lower bound on the temporal gradient for the activation of *X*_*f*_ to occur. The parameter *p* is the multiplicative factor that controls the steepness of the sigmoidal function. The second term describes the decay of *X*_*f*_ following first-order kinetics.

We simulated the model shown in Figure 3-a with the above expression for the dynamics of *X* for the three different wave periods. For the threshold parameter *θ* we chose a value such that *X* only gets activated for the short wave period (*θ* = 5 × 10^−3^ s^−1^). The parameter *k*_1_ determines the magnitude of *X* while the value of *k*_2_ determines the timescale of the decay of *X* in one cycle, and therefore of the slow increase of *X* over several cycles. For the chosen value of *θ*, the activation term of *X* is appreciably different from 0 for approximately 50 s, equating to roughly 14% of the total cycle duration for *T* = 6 min. It is then easy to determine that a value of *k*_2_ = 2. × 5 10^−3^ s^−1^ results in an increase in average CI that saturates after 6 cycles. Since the values of 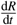 and *θ* are of the order of 1 × 10^−3^ s^−1^, *p* is chosen to be equal to 1 × 10^3^ s to make the argument of the sigmoid of the order of unity. To determine the values of *α, β*, and *γ*, we chose the simplest case of giving *M, S*, and *X* equal weightage in the definition of CI. The values *α* = *β* = *γ* = 0.15 resulted in the best fit with the experimental data. We should note that the value of *γ* is dependent on *k*_1_ which determines the amplitude of the activation of *X*.

The resulting CI for the different wave periods is shown in Figure 3-b as a function of time. Consistent with the experimental results, the CI for the short wave period shows in increase for the first several cycles and then saturates. In contrast, the CI for the long periods is unchanged for each cycle. The difference between the short and long period responses can be understood when examining the dynamics of *X*_*f*_ − *X*_*b*_ as a function of time (Figure 3, SI). For the short period waves, the time derivative of *R* exceeds the threshold value *θ*. Since the memory *M* at the front of the cell is non-zero and is zero at the back, only the front value of *X* increases. The decay of *X*_*f*_ is not rapid enough to reset it to its original value, resulting in an increase of *X*_*f*_ for the first few wave cycles, after which its mean value no longer changes. For the long periods, the time derivative does not exceed the threshold value and *X* does not accumulate appreciably. Thus, for these periods, *X*_*f*_ − *X*_*b*_ does not increase and does not contribute significantly to CI. This is also evident from the computed average CI for each cycle, shown as solid lines in Figure 3-b. The model is able to replicate both the experimentally observed slow increase in CI data as well as the cycle-independent response for long periods.

Our experimentally observed enhancement of chemotactic ability of cells due to repeated exposure to waves has clear relevance to the aggregation of *Dictyostelium*. When waves of cAMP initially arise spontaneously from random locations, they pass through the cells in all directions; the period of these waves is usually large. Over time, the wave frequency and the cAMP concentration rise [19, 36], and the center with the highest frequency or the lowest period (generally *T* ∼ 6 min), dominates the rest [14]. Then, waves with a fixed direction and frequency pass through cells. At this point, enhanced chemotaxis under periodic stimulation would help cells to reach the wave center sooner. This, combined with our earlier observations that the chemotactic ability improves as the background cAMP concentration increases [30], then leads to more optimal aggregation and, thus, better chances of survival.

Our proposed model has modules for both spatial and temporal gradient sensing that operate in parallel and independently. Temporal gradient sensing is the determinant player for bacteria, which are too small and move too fast to employ spatial sensing [38]. Since eukaryotes are large enough to sense and respond to spatial gradients [39, 40], most studies have focused on their response to spatial gradients. Our study uniquely provides evidence for the dependence of chemotaxis in *Dictyostelium* on the temporal dependence of chemoattractant gradient. These results are consistent with a recent study, which shows that migrating myeloid cells can also sense temporal dynamics of chemoattractant concentrations [41].

Our temporal sensing is formulated in terms of abstract variables, without specific identification of biochemical components. Such identification is challenging since a large number of components play a role in chemotaxis [42, 43] and additional studies are required to determine the exact biochemical components. Furthermore, we have chosen the simplest functional forms for activation thresholding and decay kinetics in Equation 2. More elaborate schemes may be possible and future work is needed to investigate this. Finally, it would be possible to further probe the temporal sensing module experimentally. Specifically, if the width of the Gaussian wave profile is decreased, keeping all else equal, the magnitude of the temporal gradient increases. Our model predicts that for these narrower profiles the CI will increase as a function of cycle number more rapidly for the short wave periods and may even increase for longer wave periods.

A.K. thanks Mahesh Mulimani for helpful discussions, and Timothy Tyree for help with software on cell-tracking.

## Supplementary Information

### I. EXPERIMENTAL SETUP

We used cells of the axenic *Dictyostelium discoideum* strain AX4, transformed with LimE-GFP (delta coil LimE-GFP) and Coronin-RFP (LimE GFP/corA RFP). Cells were grown in submerged shaking culture in HL5 medium (35.5 g HL5 powder (Formedium, Norfolk, UK) and 10 ml penicillin-streptomycin (10,000 U/ml; Gibco, Thermo Fisher Scientific, USA) per liter of DI water. When cells reached their exponential growth phase (3 − 4 × 10^6^ cells/mL), they were harvested by centrifugation at 3000 rpm for 5 minutes, resuspended in KN_2_/Ca^2+^ buffer (14.6 mM KH_2_PO_4_, 5.4 mM Na_2_HPO_4_, 100 μM CaCl_2_, pH 6.4), and washed once with the same buffer. The cells were resuspended in KN_2_/Ca^2+^ at 10^7^ cells/mL and developed for 4 and 5 hours with pulses of 50 nM cAMP added every 6 minutes.

We exposed cells to repeated waves of cAMP and recorded their movement. We performed the experiments at three different wave speeds, corresponding to a wave period of 6, 15, and 20 minutes. The microfludic wave generator is identical to the one used in an earlier study [1].

Differential Interference Contrast (DIC) images were taken every 15 seconds (for 6 minute waves) or 30 seconds (for 15 and 20 minute waves) in four fields of view spanning the width of the chemotaxis channel, 2800 μm away from the cAMP inlet, on a spinning-disk confocal Zeiss Axio Observer inverted microscope using a 10X objective and a Roper Cascade QuantEM 512SC camera. Images were captured by using Slidebook 6 (Intelligent Imaging Innovations).

Cells were plated on a glass substrate that was micropatterned with ∼ 1.5 μm thick stripes of cell adhesion-blocking polyethylene glycol (PEG) gel, as detailed in Ref. [2]. The pattern consists of 4 narrow (∼ 10 μm) and 1 wide (∼ 25 μm) untreated glass stripes separated by 30 μm wide non-adhesive PEG-gel stripes. These substrates limit the adhesion and migration of *Dictyostelium* cells to ∼ 6 − 25 μm wide stripes of non-PEG treated glass oriented in the x-direction, along the gradient and perpendicular to the flow. Thus, cell migration was effectively one-dimensional (1D), either up or down the gradient (positive or negative x-direction). This greatly simplified the collection and analysis of data as compared to 2D chemotaxis on a standard glass substrate.

### II. CELL AND WAVE TRACKING

The image analysis pipeline from DIC movies to computing chemotactic indices for the tracked cells is as follows: For each image in the movie, we applied the Scharr filter from Python’s Scikit-Image package. The filtered image is further smoothened using Gaussian blurring and was binarized to separate the cells from the background. This method of pre-processing DIC images is similar to that used in Ref. [3]. To segment the cells, we started at a pixel labeled as a cell but not segmented thus far and built a region around it by recursively including other hitherto uncounted cell-pixels that were contiguous with the current region. The region stopped growing once all contiguous pixels were identified. This process was repeated until all the cell-pixels are counted and numbered. To avoid objects that were too small or too large, we chose a lower and upper cutoff for the area of individual and isolated cells, based on a histogram of areas of all identified cells.

In order to construct tracks of individual cells, the region of a given cell in the current frame is separately used as a filter on the next frame. The region in the next frame with the highest overlap is considered to correspond to the given cell. Among the constructed cell tracks, only tracks that last at least two periods and at least three quarters of the duration of each period were considered for the next step. This was done so that we tracked the same set of cells over a prolonged period of time. Furthermore, if at any point in the track, a cell gets closer than a cutoff distance of 12 μm to another cell, that time point is excluded from the track. This is done to preclude any effects of cAMP produced by neighboring cells on chemotaxis. Once the cell tracks were constructed and selected, the chemotactic index (CI) of each cell as a function of time was calculated using Equation 1 in the main text. The average CI of the population for a given wave was then computed as the area under the CI-time curve divided by the number of bins, averaged over all the tracks in that cycle.

To visualize the cAMP wave, a fluorescent dye, Alexa Fluor 594 Hydrazide (Invitrogen), was added to the central stream along with cAMP at a concentration of 1000 nM. The fluorescence intensity in each frame was fitted to a univariate Gaussian profile 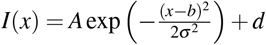 where *x* is the distance from the peak of the wave along direction of propagation, and *A, b, σ, d* are, respectively, the amplitude, position of the peak, width of the Gaussian, and the background intensity. *I*(*x*) is the intensity along the direction of the propagation after averaging in the perpendicular direction. While simulating the wave, we used the function

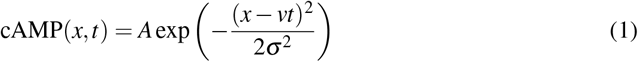

where *A* = 1000 nM, *v* = 1400 μm*/T* where *T* is the wave period in seconds (1400 μm is the distance a wave moves in each period from start to finish), *σ* = 120 μm.

### III. TRENDS IN CHEMOTACTIC INDEX

The mean value of Chemotactic Index (defined in Equation 1 in the main text) in a cycle increases upon exposure to multiple waves if the wave period *T* is short and remains low and unchanged if *T* is large. This is due to an increase in the directionality of cells when *T* is small; the (undirected) cell speed remains roughly the same (∼ 3 − 4 μmmin^−1^) for all cycles and periods, as shown in Figure 1.

When the chemotaxis experiments were repeated with cells that were developed for four hours, the cells still showed a significant increase in the mean chemotactic index for the *T* = 6 min waves, similar to the cells developed for five hours, suggesting that the increase in the chemotactic ability is in response to exposure to the chemoattractant waves, and not merely the passage of time. See Figure 2.

### IV. MODEL DESCRIPTION

The complete model used is an extension of the LEGI+M model along with Equation 2 of the main text. We used the LEGI+M model as detailed in an earlier study [4]. The entire set of equations and the parameters are reproduced here for completeness.

**FIG. 1:**
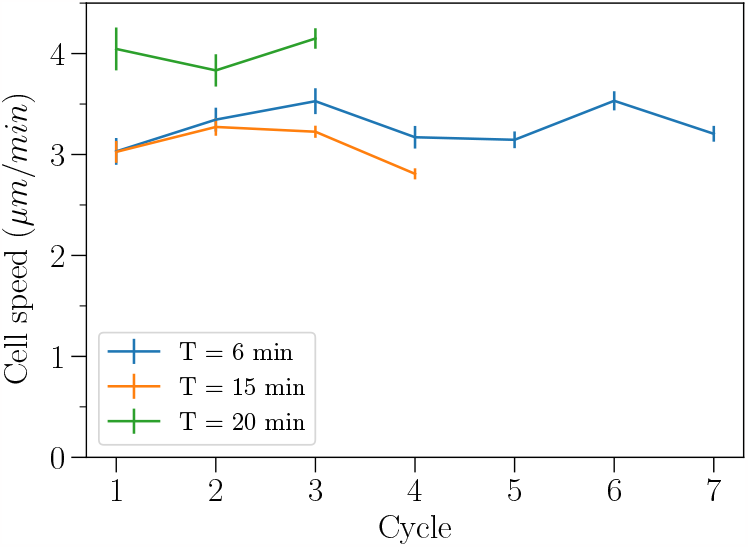
The average undirected cell speeds remain roughly the same for all cycles and wave periods. Error bars in the plot denote the standard error of the mean.

**FIG. 2:**
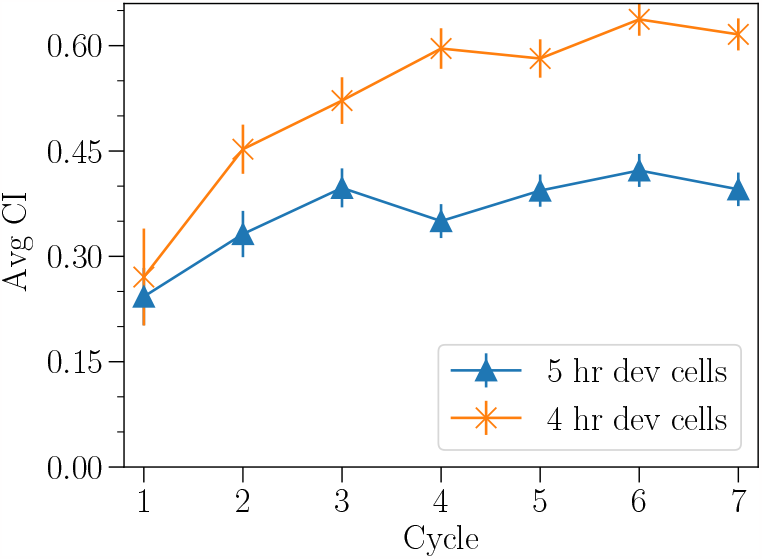
Average CI increases as a function of cycle number when cells were developed for four or five hours prior to the exposure of cAMP waves.

External cAMP activates the receptors on the surface of the cell. Two kinds of receptors *R*_1_ and *R*_2_, differing in their binding affinities and constitutive activations are considered. The two types of receptors have the same downstream effect, so their sum *R* = *R*_1_ + *R*_2_ is considered in the rest of the model. *R* activates *E* and *I*, which respectively excite and inhibit the response element *S* following zeroth-order ultrasensitivity kinetics, so that a weak gradient in *R* and *E* is amplified to a strong gradient in *S. S* also promotes the switching of the bistable memory module *M* from 0 (low) to 1 (high), which feeds back to activate *S*, particularly on the back half of the wave. The new component *X* is activated when *M* is high and when the temporal gradient of *R* is above a positive threshold *θ*. All the components of the model except *I* are localized at the membrane. Thus they take two values, one at the front and one at the back, denoted by the subscripts *f* and *b* respectively. Equation 2 shows the equations governing the kinetics of (global) *I*, the model components at the front of the cell, and the definition of the chemotactic index (CI). A corresponding set of equations exists for the components at the back of the cell.

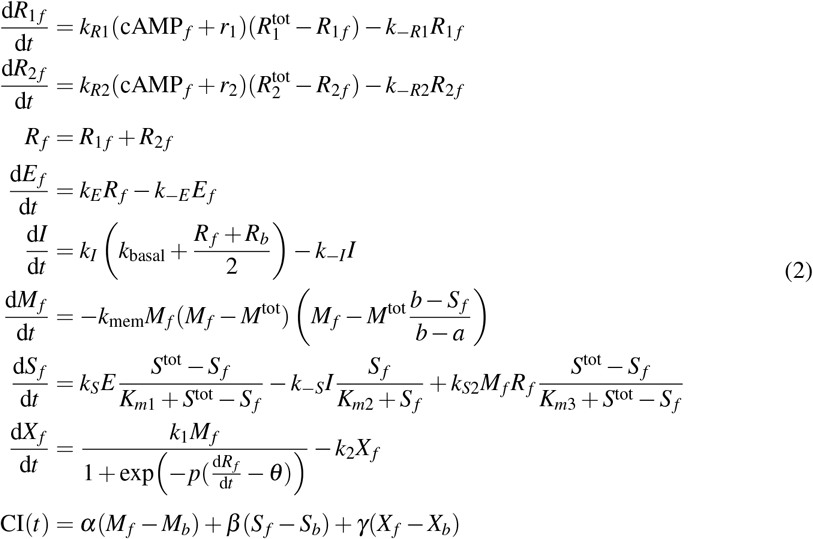

The parameters of the model are shown in Table I. The concentration of cAMP is measured in nM and the rest of the components are de-dimensionlized or unitless.

The model equations are integrated using MATLAB’s ODE solver ode23tb [5], which is an adaptive Runge-Kutta method with a variable step size. The cell is assumed to be one-dimensional with a length of *L* = 10 μm. The cAMP stimulus is provided according to Equation 1. The center of the cell is assumed to be at *x* = 0 and *t* ranges from −*T/*2 to *T/*2 for each cycle. To prevent *M* from getting stuck at the extrema *M* = 0 and *M* = 1, a “kick” is given to *M* every 15 s in the simulation: *M* is set to 0.05 if *M* = 0 and to 0.95 if *M* = 1.

#### A. Spatial versus Temporal Sensing

We have argued in the main text, along with Figure 2 therein, why a model for the activation of *X* that depends on the *concentration* of cAMP or any other component downstream of it fails to reproduce the experimental trend in ⟨CI⟩. Here we explicitly show the results of integrating a model where the rate equation for *X* obeys simple mass action kinetics dependent on concentrations. Consider the model with an equation for *X*_*f*_ given by

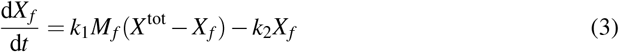

with a similar equation for *X*_*b*_. Equation 2 is integrated along with this equation, and the resulting time series of *X*_*f*_ − *X*_*b*_ is plotted as a function of time in Figure 3 (top panel). The levels of *X*_*f*_ − *X*_*b*_ for long periods exceed that for the short period. As explained in the main text, this happens because in the case of long periods, the cells perceive high levels of *M* for a prolonged amount of time, and that leads to stronger activation. This is contrary to our purpose of introducing *X* as a means to explain rising ⟨CI⟩ for short period only. On the other hand, the temporal model shows, in Figure 3 (bottom panel), that the values of *X*_*f*_ − *X*_*b*_ stay higher for the short period than for the long periods.

**FIG. 3:**
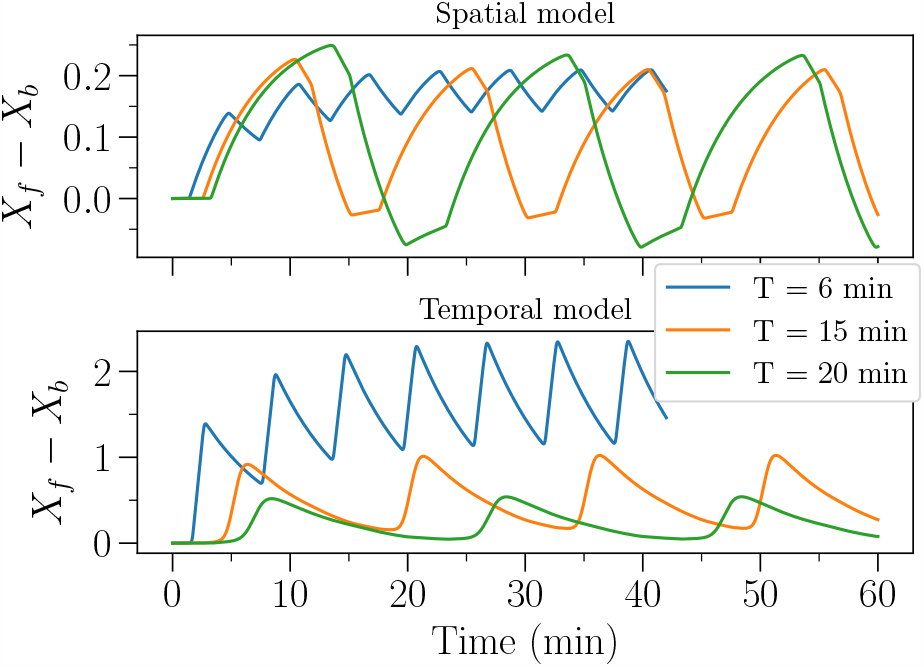
Comparison of time series of *X*_*f*_ − *X*_*b*_ when the kinetics of *X* is given by equation 3 of the spatial model (above) and the temporal model (below). For the spatial model, *k*_1_ = 1.0 × 10^−3^ s^−1^, *k*_2_ = 2.5 × 10^−3^ s^−1^, and *X* ^tot^ = 1. The parameters for the temporal model are as defined in Table I.

## Notes

### Competing Interest Statement

The authors have declared no competing interest.

